# A simple, inexpensive and multi-scale 3-D fluorescent test sample for optical sectioning microscopies

**DOI:** 10.1101/2021.04.27.441588

**Authors:** Ilya Olevsko, Kaitlin Szederkenyi, Jennifer Corridon, Aaron Au, Brigitte Delhomme, Thierry Bastien, Julien Fernandes, Christopher Yip, Martin Oheim, Adi Salomon

## Abstract

Fluorescence standards allow for quality control and for the comparison of data sets across instruments and laboratories in applications of quantitative fluorescence. For example, users of microscopy core facilities expect a homogenous and time-invariant illumination and a uniform detection sensitivity, which are prerequisites for quantitative imaging analysis, particle tracking or fluorometric pH or Ca^2+^-concentration measurements. Similarly, confirming the three-dimensional (3-D) resolution of optical sectioning micro-scopes prior to volumetric reconstructions calls for a regular calibration with a standardised point source. Typically, the test samples required for such calibration measurements are different ones, and they depend much on the very microscope technique used. Also, the ever-increasing choice among these techniques increases the demand for comparison and metrology across instruments. Here, we advocate and demonstrate the multiple uses of a surprisingly versatile and simple 3-D test sample that can complement existing and much more expensive calibration samples: simple commercial tissue paper labelled with a fluorescent highlighter pen. We provide relevant sample characteristics and show examples ranging from the sub-µm to cm scale, acquired on epifluorescence, confocal, image scanning, two-photon (2P) and light-sheet microscopes.

**Graphical abstract:** Pyranine-labeled tissue paper, imaged upon 405-nm epifluorescence excitation through a 455LP LP dichroic and 465LP emission filter. Objective ×20/NA0.25. Overlaid are the normalised absorbance (dashed) and emission spectra (through line), respectively. In the present work we show that this “primitive” and inexpensive three-dimensional (3-D) test sample is a surprisingly versatile and powerful tool for quality assessment, comparison across microscopes as well as routine metrology for optical sectioning techniques, both for research labs and imaging core facilities.

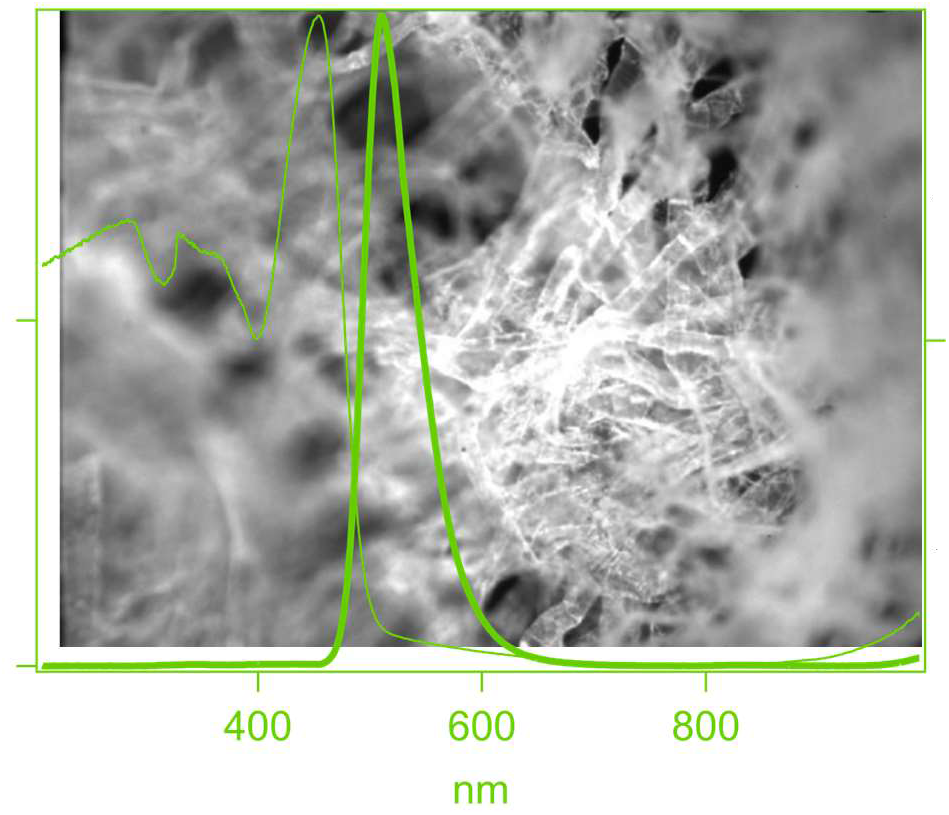

**Research highlights:**

- highlighter-pen marked tissue paper is a surprisingly powerful and versatile test sample for 3-D fluorescence microscopies
- standard tissue paper presents features ranging from 400 nm to centimetres
- our sample can simultaneously be used for testing intensity, field homogeneity, resolution, optical sectioning and image contrast
- it is easy to prepare, versatile, photostable and inexpensive

## INTRODUCTION

Fluorescence microscopy is indispensable for interrogating spatial and temporal relationships among structures on surfaces or in volumes, not only in medicine and biology, but also in the material sciences and engineering disciplines. Often, the objects under study are neither flat nor two dimensional. As a consequence of their limited numerical aperture (NA) and the associated “missing cone” problem, optical microscopes, have a poorer capacity to discriminate structures in axial (*z*) than in lateral directions (*xy*), within the focal plane. Also, three-dimensional (3-D) samples are often labelled throughout their volume, and images will contain out-of-focus information from fluorophores above and below the focal plane. In this context 3-D imaging requires either the serial slicing of the sample into thin sections or else the use of optical sectioning that keep the sample intact. 3-D optical sectioning techniques rely either on excitation confinement, **Fig. 1***A* (like two-photon excitation fluorescence (2P) microscopy, *left*, or light-sheet microscopies, *middle*), or they are based on emission spatial filtering (e.g., fluorescence detection through a confocal pinhole, *right*). Sometimes such approaches are combined with post-acquisition image processing, like exhaustive photon reassignment or ‘deconvolution’ techniques for improving image contrast, see **Box1**.

**FIGURE 1.**
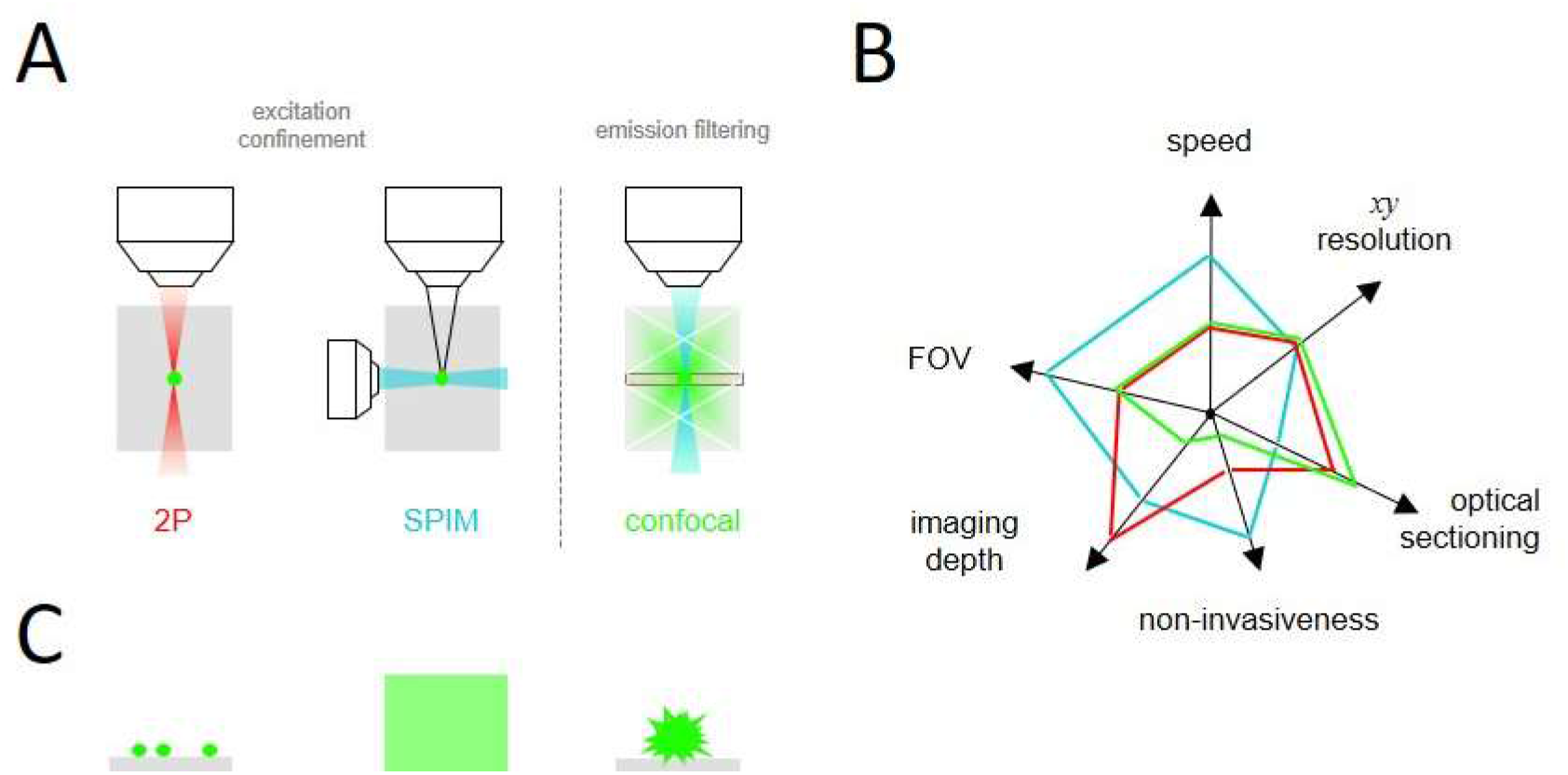
Optical sectioning microscopies and available 3-D test samples. (A), 3-D microscopic imaging relies either on the confinement of fluorescence excitation, as in the case of two-photon (2P) and selective-plane illumination microscopy (SPIM), or emission spatial filtering, as in confocal detection, where only the fluorescence emitted in the focal plane passes a tiny pinhole and reaches the detector. (B), different optical sectioning techniques excel in different image parameters. Colour code as in panel A. FOV - field of view. (C), common test samples. *Left -* fluorescently labelled microspheres, *middle* - homogenous test slides or dye solutions, *right* - natural autofluorescent pollen grain.

Optical sections can be of the order of a micrometer (µm) for deconvolution microscopies, confocal laser-scanning microscopies (CLSM) and 2P microscopies (Denk, Strickler et al. 1990, Diaspro 2002, Pawley 2006). On the other extreme, light-sheet microscopes typically discriminate axial features of tens of µm, with an even lower axial sectioning achieved on fluorescence macroscopes (Huber, Keller et al. 2001) that use long-working distance air lenses with moderate numerical apertures (NA). In fact, the different ultra-microscopes (Siedentopf and Zsigmondy 1902, Dodt, Leischner et al. 2007) - see (Masters 2020) for a recent comment -, orthogonal-illumination (Voie, Burns et al. 1993) or selective-plane illumination microscopes (Huisken, Swoger et al. 2004) form a heterogeneous family of techniques that all feature a 90°-angle between the illumination and detection optical paths. They are now collectively called light-sheet microscopes, see (Keller and Dodt 2012) for a historical perspective.

Despite of the different spatial scales probed by all these fluorescence microscopies, their different fields of view (FOV) and different speeds of acquisition as well as variable degrees of light-exposure to the sample (see **Fig. 1***B*) the user faces a similar challenge: he or she must evaluate, which microscope responds best to the question under study, and then test and calibrate the instrument against a test sample of known dimensions to interpret the 3-D image. Such a test sample should present a stereotyped 3-D structure and it should offer some flexibility in terms of excitation and emission wavelengths. It should be resistant to photobleaching and either it should have a long shelf life or be cheap and easy to prepare freshly and in a reproducible manner. To be widely applicable and allow comparison across different microscopy techniques it would be advantageous if the test sample would offer features spanning different spatial scales.

*What are the available commercial fluorescence standards?* By and large, they fall into point sources, samples presenting some structural features and homogeneous, uniformly labelled samples, **Fig. 1***C*. Tiny fluorescent beads come in many colours and sizes, and sub-diffraction dye-labelled polystyrene microspheres are a de-facto standard as fluorescent point sources (Resch-Genger, Hoffmann et al. 2005). Bigger, µm-sized beads have serve as a reliable intensity standard, e.g. for quantitative Ca^2+^ imaging (Neher 1995), but, unfortunately, the use of ‘bead units’ (BU) for quantitative fluorimetry never really took off. Microspheres are fairly expensive and they tend to aggregate during storage, too. Also, the sub-100-nm beads are quite dim, making their use in thick 3-D samples difficult. Also, whereas bead monolayers on a glass coverslip are produced without much effort, embedding of beads in agarose gels to produce homogenous ‘raisin cake’ 3-D samples is cumbersome and work-intensive (Oheim, Salomon et al. 2019). On the other hand, uniform thick test samples like the popular Chroma fluorescent plastic slides, lack distinctive features other than the surface to focus at, and they are too absorbing and optically turbid for deep imaging with techniques other than nonlinear microscopy, **Fig. 1***C* (*middle*). Pollen grain has emerged a popular test for instrument performance (Potter 1996, Vitha, Bryant et al. 2009, Kirkby, Nadella et al. 2010, Siegel and Brooker 2014, Thériault, Cottet et al. 2014, Kim, Lee et al. 2018, Shin, Kim et al. 2018, Sivaguru, Urban et al. 2018, Rakotoson, Delhomme et al. 2019), **Fig. 1***C* (*right*). Featuring broad excitation and emission spectra and - at least for the thorny-type variant a characteristic ‘sea mine’ aspect – naturally autofluorescent pollen grains combine structural features in the 1- to 10-µm range that make them perfect 3-D objects for confocal and 2PEF laser-scanning microscopes. Yet, they are too small for mesoscopic imaging techniques like the ever-growing family of light-sheet microscopes (Hillman, Voleti et al. 2019, Wan, McDole et al. 2019) that sample cubic volumes with hundreds of µm up to mm side lengths. More recently, Argolight test samples offer a more complete but expensive solution for multi-parametric metrology (Royon and Converset 2017). These commercial test samples contain several fluorescent patterns. Here, each pattern is designed to assess one or several parameters of a microscope: resolution, field uniformity, intensity response, co-registration accuracy between channels etc.. Nevertheless, a broadly available, inexpensive and versatile test sample spanning the µm to cm range has been missing. Also, in practical applications, the aim is often not a rigorous calibration and quantification but a “quick and dirty” verification of the microscope performance before or during an experiment. Such a sample for troubleshooting and on-the-fly diagnosis must be above all readily available and easy to use.

In the current paper, we present and characterise such a 3-D test sample that can complement existing protocols and calibration routines on a daily basis. It requires only three components readily available in any research laboratory: standard tissue paper (e.g., KimWipe^®^), a yellow highlighter pen (e.g., Textmarker, Stabilo Boss^®^,…), and, for light-sheet imaging, a plastic mount produced on a standard 3-D printer. We provide 3-D images of structural features ranging from sub-µm dimensions to the cm scale. Examples images acquired on epifluorescence, confocal, image scanning (Müller and Enderlein 2010) (Airyscan), 2P and light-sheet fluorescence microscopes are provided. Our results also confirm the interest of a new 2P-spinning disk microscope (Rakotoson, Delhomme et al. 2019) for highly resolved large-field 3-D imaging.

## MATERIALS AND METHODS

### Calibration sample preparation

Small stripes of tissue paper (Kimberly-Clark, Irving, TX) were fluorescently labelled with a yellow highlighter pen (Stabilo Boss^®^, Schwan-Stabilo, Weißenburg, Germany) in a single swipe, mounted on a microscope slide and covered with a #1.5 cover glass (Marienfeld Superior, ThermoFisher) that was held in place with tiny stripes of adhesive tape, **Fig. 2***A*. For the light-sheet microscopies, we attached the dye-stained paper to a custom 3-D printed plastic holder that maintained the paper at a 45°-angle in a small side-open cube, similar to a fluorescence filter cube.

**FIGURE 2.**
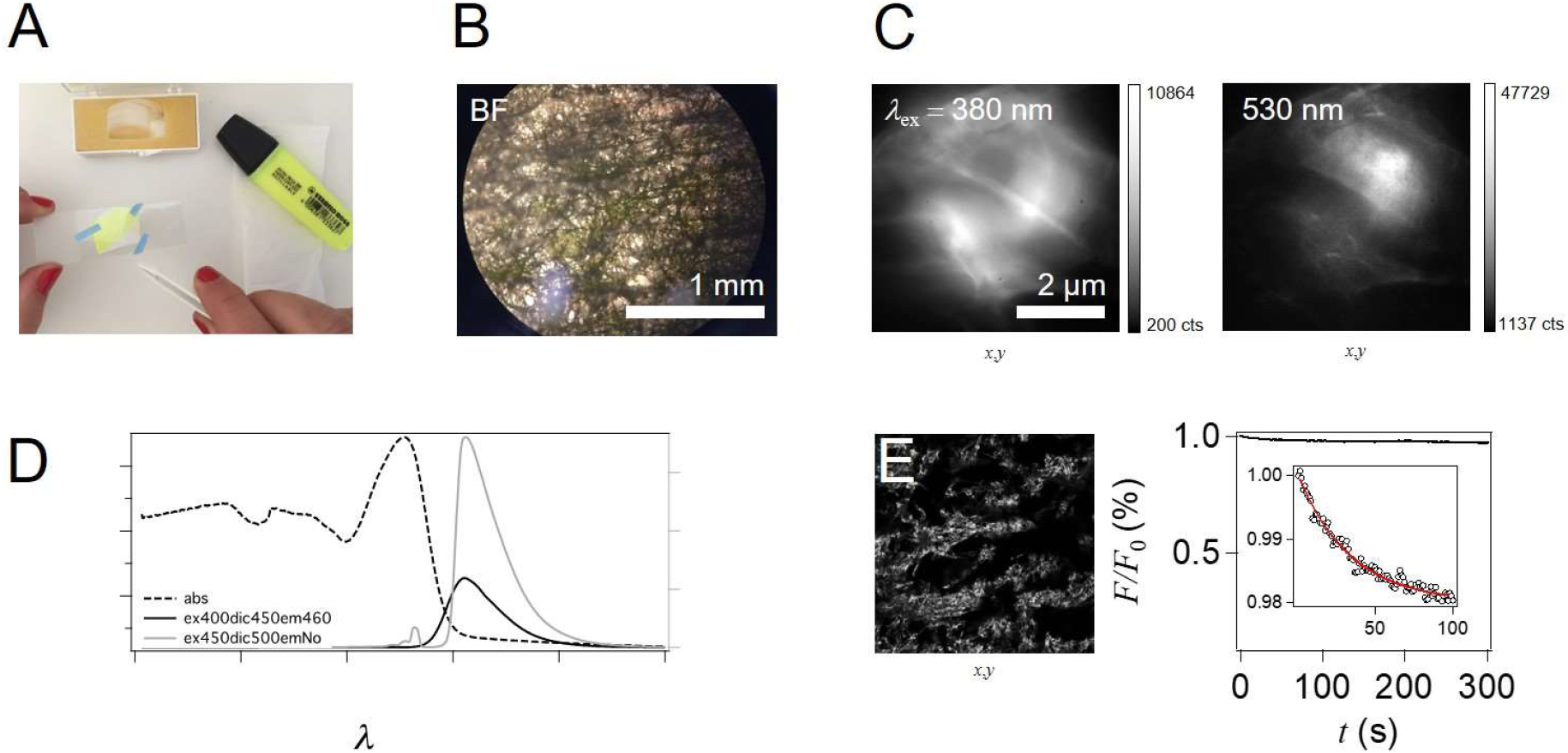
Fluorescently labelled tissue paper is a versatile test sample. (A), sample preparation. The standard tissue paper was fluorescently marked with a highlighter pen and placed between slide and coverglass, the later was held in place with small stripes of scotch. (B), transmission view with a ×10/0.3air objective. Scale bar, 1 mm. (C), higher-magnification epifluorescence images upon UV and green-light excitation. Scale bar, 2 µm. (D), measured absorbance (dashed) and emission (solid) spectra upon 400-nm (*black*) and 450-nm excitation (*grey*), respectively. (E), *left*, confocal micrograph of a single plane of fluorescently labelled tissue paper and, *right*, evolution of fluorescence with time upon continuous excitation. *Inset* zooms in on the time course of the 2% bleaching component.

### Imaging spectroscopy

Transmission, absorption and fluorescence emission spectra were measured on an imaging microspectrometer, combining an inverted microscope (IX83, Olympus) equipped with a ×20/NA0.25 air objective, an IsoPlane SCT320 spectrometer and a PIXIS 1024 eXcelon^™^ electron-multiplying charge-coupled device (EMCDD) camera (TeleDyn Princeton Instruments). We used a 600-nm blaze/50 grooves/nm grating, corresponding to a 0.77-nm spectral resolution. For transmission and absorbance measure-ments acquisition parameters were *t*_exp_ = 300 ms, slit width = 150 μm, minimal iris, aperture 20X, all controlled via µMAN-AGER software (https://micro-manager.org). For fluorescence acquisition the exposure time, *t*_exp_, was 50 ms, and lamp intensity 100%, all other things equal. Fluorescence was excited using a Xe-arc lamp at either 405 nm or 450 nm and imaged through a 455LP (505LP) dichroic with 465LP (no) emission filter, respectively. Samples were either a clean 2-mm thick naked microscope slide for background subtraction or the slide carrying a highlighter-coloured tissue paper, as described above. Transmission spectra were corrected using the bare glass slide as a reference. Absorbance was calculated as *Abs* = 2-log_10_(*T*), where *T* is the measured transmission in percent. Fluorescence spectra are reported in arbitrary units (AU) after background subtraction. Spectra are averages over 1023 pixel rows. The image in the graphical abstract was acquired on a small CMOS camera mounted on the imaging arm of the same microscope.

### Epifluorescence

Wide-field fluorescence images were acquired on a home-built multi-modal microscope (van’t Hoff, Reuter et al. 2009). Briefly, the quasi monochromatic (Δλ = 18 nm, FWHM) output of a polychromatic light-source (Poly II, TILL Photonics, Gräfelfing, Germany) was coupled via a quartz fibre to a breadboard upright microscope, via a 40-mm achromatic converging lens, a 45°-filter cube and ×63/NA0.8w dipping lens (Zeiss, Oberkochen, Germany). Fluorescence was excited at 380 nm or 530 nm, either using a Fura-2 cube (F-76-520, AHF Analysentechnik, Tübingen, Germany, see **Supplementary Table S1** in the **Supporting Material Online**) or a TRITC filter cube (F36-503), respectively. Fluorescence was imaged through an Olympus tube lens assembly (*f*_TL_ = 180 mm) onto an EMCCD camera (Cascade512B, Roperscientific, Parray Vielle Poste, France), after ×2 additional magnification. The pixel size in the sample plane was 109 nm/px at a ×137 effective transverse magnification. For acquisition and analysis we used METAMORPH software (Molecular Devices, San José, CA). Images in Fig. S2 were taken at ×63/NA0.25 air with or without additional ×2 magnification.

### Confocal microscopy

Confocal micrographs in Fig. 2 were taken on a standard ZEISS LSM710META microscope upon 488-nm excitation, using a 488LP beamsplitter, and variable emission spectral windowing on the META detector. With a ×10/0.3NA air objective (ZEISS EC Plan-Neofluar) the pixel size was 1.58 µm/px. The 1-Airy diameter of the confocal pinhole corresponded to a 13-µm sectioning.

### Image scanning microscopy

We used image scanning microscopy (Müller and Enderlein 2010) on a LSM880 Airyscan inverted microscope with a ×63/1.4NA oil-immersion objective (ZEISS Plan-Apochromat M27) for the acquisition of the images shown in Fig. 3 and Movie S1. The pinhole was fully open and the 32-element ‘compound-eye’ detector was used instead to attain an effective 200- to 210-nm lateral (*xy*) and 510-nm axial (*z*-) FWHM resolution, measured with 93-nm beads. We did not apply any correction for the finite bead size (Nadrigny, Rivals et al. 2006, Zhang, Zerubia et al. 2007, Barentine, Schroeder et al. 2018). The pixel size in the sample plane was 89 nm/px, optical sections were taken at 0.3 µm spacing. Fluorescence was excited using the 488-nm line (at 15% power) and detected beyond 495 nm. We systematically scanned with a line-averaging of four.

**FIGURE 3.**
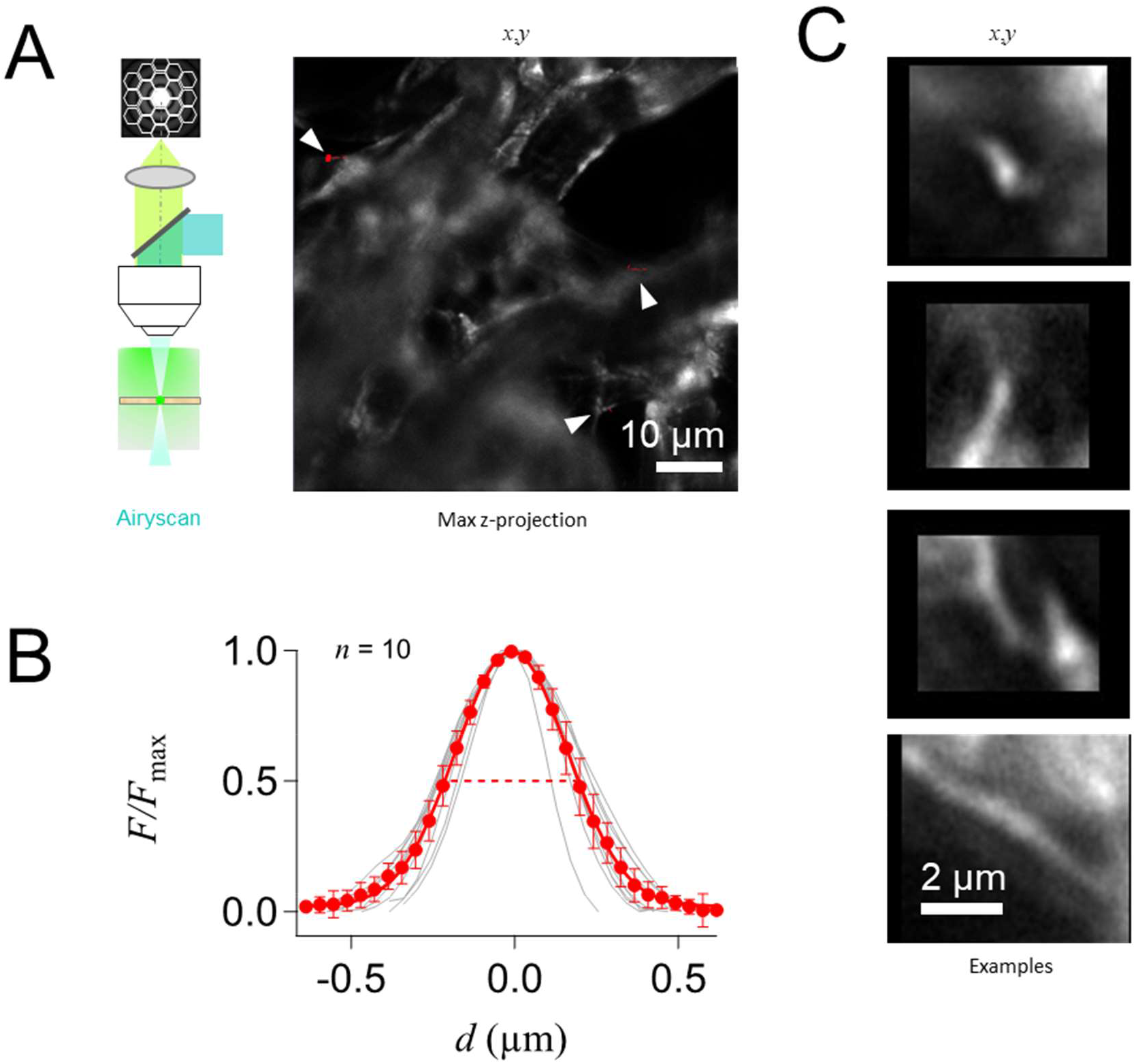
Pyranine-labelled tissue paper contains sub-wavelength features. (A), *left*, schematic optical layout of the used Airyscan confocal microscope that uses a ‘honeycomb’ multi-pixel detector instead of a confocal pinhole and single photomultiplier. *Right*, maximum-intensity projection of a *z*-stack of 488-nm excited fluorescence images. Optical sections were taken at 0.3 µm spacing. (B), examples of sub-wavelength features identified in (C) the upper image volume, *left*, and normalised intensity line-profiles of tiny features (*grey)* and their ensemble average and SD (*red*), *right*. Measured FWHM (*dashed*) was 404 ± 3 nm (*n* = 10).

### 2P-spinning-disk microscopy

Non-linear fluorescence images in Fig. 4 were acquired upon fs-pulsed 820-nm centre-wavelength (CWL) excitation on a custom 2P-planar illumination microscope (Rakotoson, Delhomme et al. 2019). The expanded and attenuated output of a Titanium-Sapphire laser (MaiTai HP, Spectra Physics, Palo Alto, CA) with external prism compressor (‘DeepSee^™^’) was injected into the microscope, and ∼50 excitation spots simultaneously scanned across the field of view. The fast multi-spot scanning of the OASIS microscope results from a unique modified Nipkov-Petráň-type spinning-disk geometry, in which 5,000 microlenses and their corresponding confocal pinholes (realised as the holes of a perforated 715LP dichroic coating) are combined on the same custom-designed and -manufactured single disk. The detection is mildly confocal (2 Airy), and the disk spins at a constant speed of 5,000 RPM, corresponding to a Nyquist-limited exposure time of 6 ms, full frame. All exposure times were integral multiples of this. The resulting fluorescence was short-pass filtered (**Table S1**) and the ‘green’ (400-565 nm) colour channel detected on half of the chip of a scientific complementary metal-oxide semiconductor (sCMOS) camera (PCO.edge4.2, Kelheim, Germany). The OASIS microscope was controlled through SIAM software (TILL.id, Planegg/Martinsried, Germany). Integration times were 12 and 48 ms, respectively, for images taken with the ×20/NA0.8air and ×25/NA1.1w objective (Nikon, Champigny-sur-Marne, France), pixel sizes in the sample plane were 182 (228) nm for the dipping (air) objective.

**FIGURE 4.**
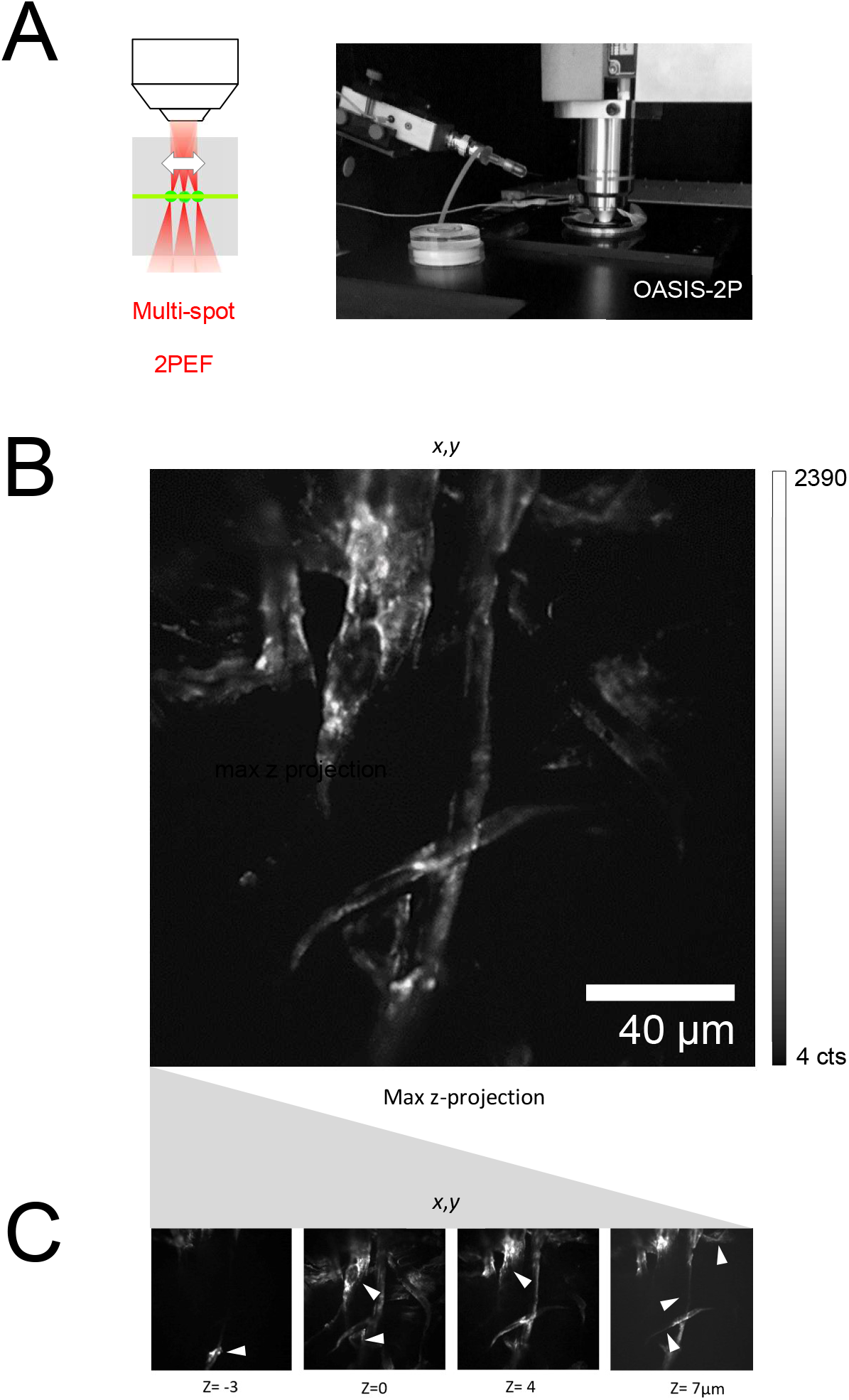
2P-imaging of pyranine-labelled tissue paper. (A), *left*, schematic illustration of the used multi-spot scanning, resulting in planar non-linear fluorescence excitation, and photo of the nosepiece of the upright, home-built OASIS (<u>O</u>n-<u>A</u>xis 2-Photon Light-<u>S</u>heet Imaging <u>S</u>ystem) microscope. Note the clear headroom around the recording site. Objective was a ×20/NA0.8air lens. (B), maximum-intensity projection of a *z*-stack of 2P(820-nm) excited fluorescence images taken at 1-µm axial distance, and example planes, (C). Arrowheads identify axial features discriminated on subsequent planes.

### Selective-plane illumination microscopes (SPIM)

We used two different light-sheet microscopes. For Fig. 5 and Movie S1, we employed a home-built, modular light-sheet microscope (**Supplementary Fig. S1**), which allows for swapping the excitation optics, imaging medium and objectives. The imaging arm on the system is set up vertically to allow for the use of dipping objectives. The beam of a 488-nm Coherent Sapphire laser, was expanded 6.66-fold, shaped with a plano-convex *f* =40 mm cylindrical lens and focused by an Olympus SLMPlan ×20/NA0.35 objective to a planar, unidirectional light sheet. An identical objective lens was used on the orthogonal collection arm. The collected fluorescence was filtered and imaged onto a large-format sCMOS detector (ANDOR Neo, pixel size 6.5 µm, Oxford Instruments, Belfast, Ireland). The acquisitions were controlled with µMANAGER software. At ×20 effective transversal magnification, the pixel size in the sample plane was 390 nm/px, the step size for *z-* acquisitions was 1 µm.

**FIGURE 5.**
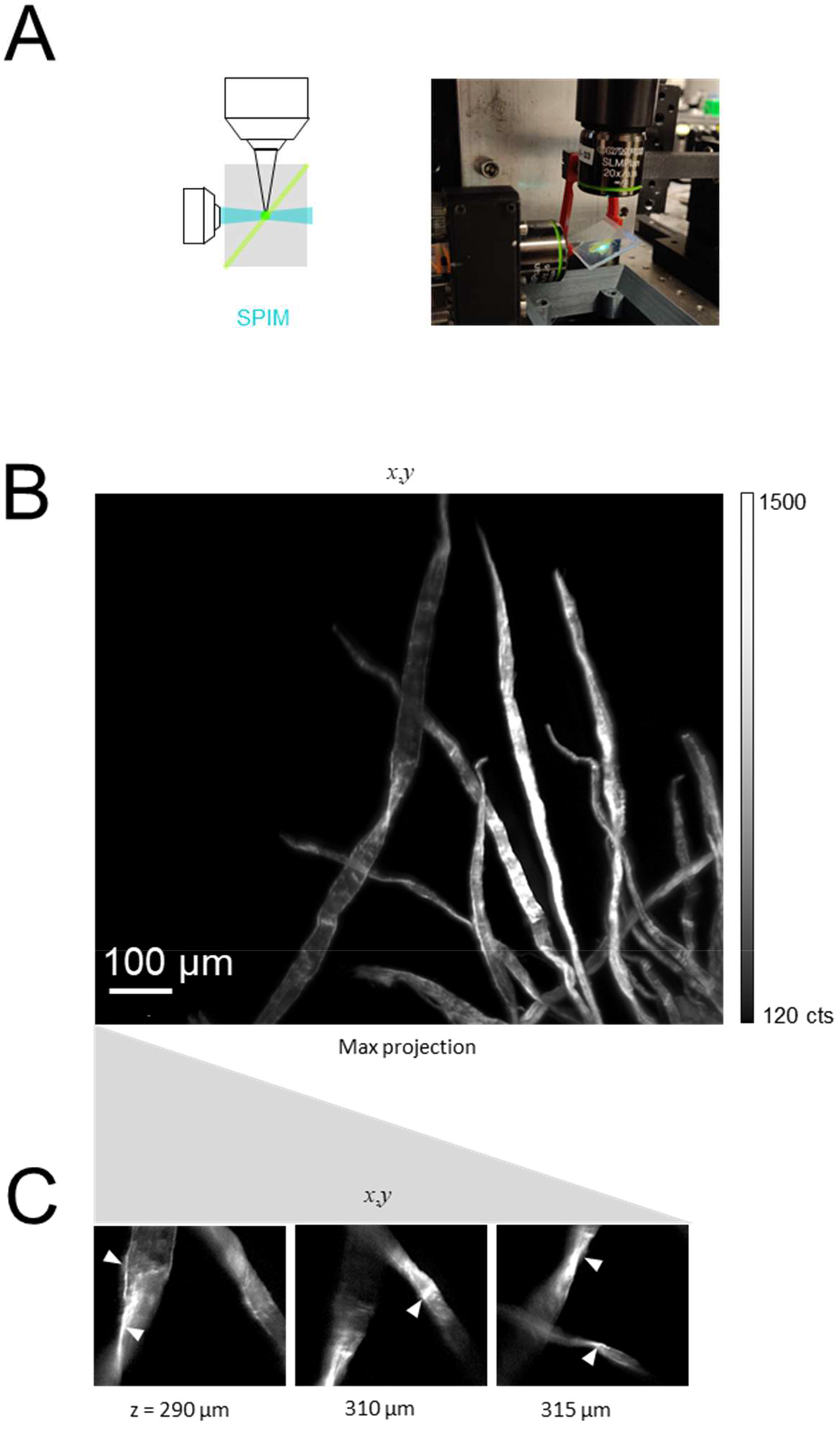
Light-sheet microscopic imaging of fluorescent tissue paper. (A), *left*, schematic optical layout of the used orthogonal excitation-imaging geometry and photo of the home-built SPIM, *right*. (B), maximum-intensity projection of the low-density edge of fluorescent tissue paper, and examples of single optical planes, (C), at identified axial positions.

The data shown in Fig. 6 and Movie S3 was acquired on a LaVision UltraMicroscope II, using a single light sheet out of the 3 crossed beams used for illumination. With an optical zoom from ×1.26 to ×12.6, this macroscope offers a large field of view (FOV, image diagonal up to 17.6 mm) and it is equipped with a white-light laser (SuperK extreme, NKT Photonics) and an ANDOR Neo sCMOS detector. Images shown were taken with a ×2 objective (Olympus MV PL APO 2XC, NA 0.5air) with a zoom of ×0.63 (⇔ ×1.26). The effective pixel size was 4.8 µm/px and the *z-*step size 2 µm.

**FIGURE 6.**
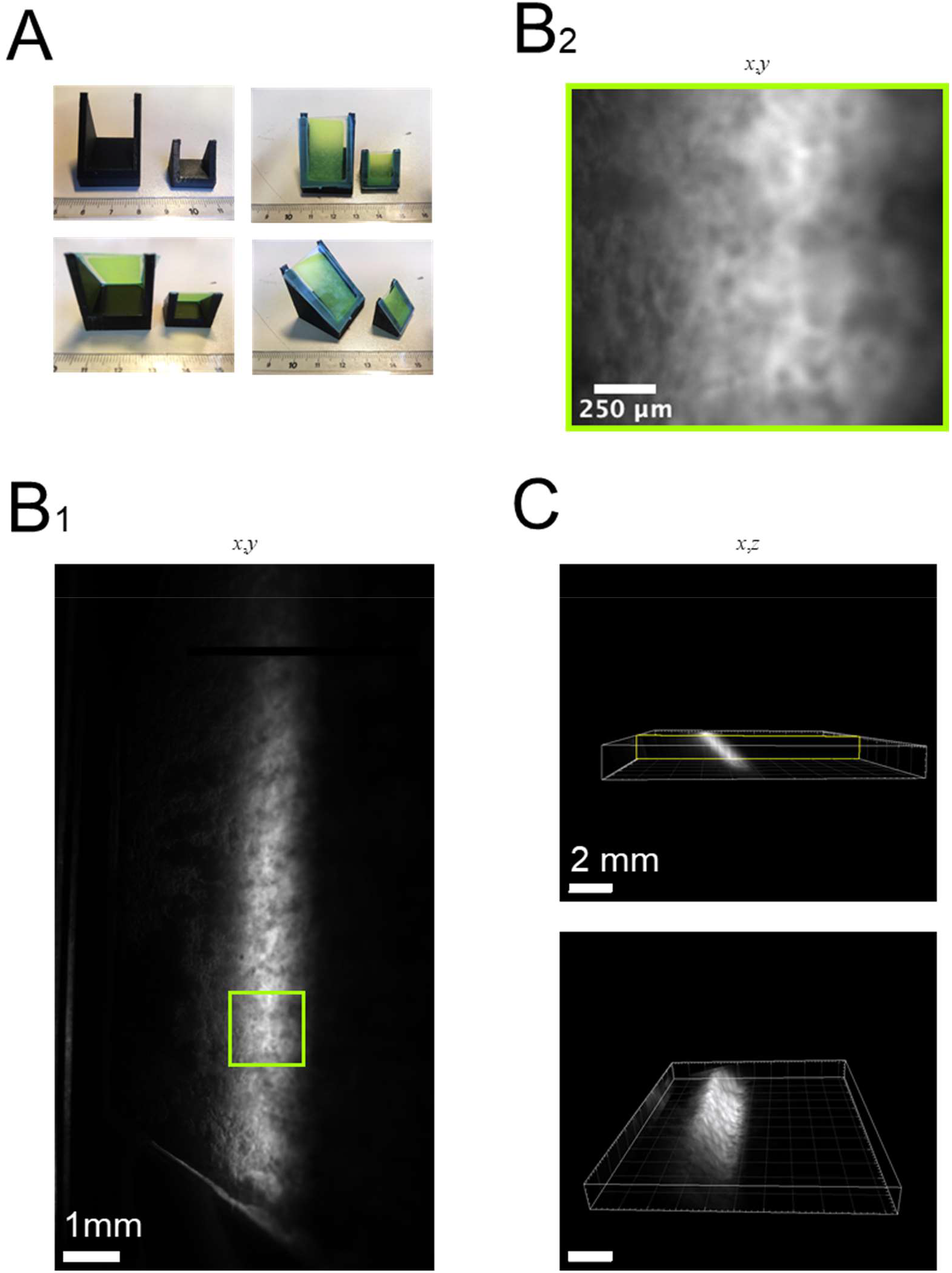
Light-sheet macroscopic imaging of fluorescent tissue paper. (A), photographs of the 3-D printed holder with and without mounted large-scale tissue paper. (B), single-plane macroscopic fluorescence image (magnification ×1.26/NA0.5, B1) and zoom on the boxed region (B2). (C), 3-D rendered view of the entire image stack shows the slanted 45°-sample geometry and a homogeneous labelling and detection over a cubic volume of (2 mm)^3^.

Filters and details for the different microscopes are compiled in **Table S1** in the **Supplementary Material Online**.

### Image processing, data analysis and statistics

Images shown are raw data, after subtraction of a dark image (i.e., with the respective light source shuttered). Intensities are reported in arbitrary units (AUs) on a grey value look-up table (LUT). No attempt was made to calibrate the respective detectors in terms of absolute photon numbers. Experiments were systematically conducted as triplicates. Reported values are average ± standard deviation (SD), of *N* independent measurements, unless otherwise stated in the figure legends. We used ImageJ64 (FIJI) (Schindelin, Arganda-Carreras et al. 2012), METAMORPH, ZEISS ZEN offline, IMARIS (Bitplane) for image acquisition, treatment and display (see figure legends). Graphs and statistics were done using IGOR PRO (Wavemetrics).

## RESULTS

### Pyranine-labelled tissue paper as a versatile test sample

Inexpensive materials readily available in the lab can replace elaborate and expensive test samples. Labelling standard bench-top tissue paper with a yellow highlighter pen, **Fig. 2***A* produced a surprisingly homogenous fluorescent sample with image dimensions limited only by the field-of-view of the respective microscope. On bright-field images, tissue paper showed a relatively uniform, fibrous, translucent aspect and axial dimensions large enough to focus on multiple layers of crossing fibres, **Fig. 2***B*. A quick check on a multispectral epifluorescence microscope revealed a broad fluorescence excitation, with a detectable signal from UV excitation at 380 nm up to green-light illumination at 530 nm, **Fig. 2***C*. The measured absorbance peaked at 452 nm with a broad shoulder in the UV. Upon 450-nm excitation, fluorescence emission (grey) extended from 470 nm to above 600 nm, with a peak detected at (512±2) nm upon off-peak excitation at 400 nm to better capture the ‘true’ shape of the spectrum (black), **Fig. 2***D*. These spectra are compatible with pyranine (8-Hydroxypyrene-1,3,6-trisulfonic acid trisodium salt, HPTS, CAS 6358-69-6, (De Borba, Amaral et al. 2000)), for which we measured very similar spectra with absorbance and emission peaks at 454 nm and 520 nm, respectively (see **Fig. S2**).

With 488-nm excitation we acquired during 5 min continuously confocal micrographs at 1Hz to study the photostability of the sample. Fitting a single exponential function with the measured fluorescence decay (*inset*), we found a very small bleaching amplitude of d*F*/*F*0(max) = 0.021 ± 0.001, meaning that 98% of the fluorescence persisted after minutes of illumination. The bleaching time-constant for this 2%-intensity loss was τ = (31.53 ± 1.84) s, **Fig. 2***E*.

Taken together, its ease of use, the molecular brightness (*B* = ε ϕ = 21,800 mol^-1^cm^-1^ 0.82 = 17,876 mol^-1^cm^-1^) (De Borba, Amaral et al. 2000, Taniguchi and Lindsey 2018), high photo stability and convenient absorbance and emission spectra make pyranine suitable for excitation with many common laser lines, and allow for a detection in the “green” to “orange” fluorescence bands characteristic for many popular small-molecule organic fluorophores and fluorescent proteins (e.g., FITC, Alexa488, TRITC, EGFP, EFYP). Thus, textmarker-labelled tissue paper is detectable with a number of filter sets routinely available in routine ‘biological’ microscopy.

### Probing optical sectioning over several orders of magnitude

Optical sectioning microscopes probe very different spatial scales, ranging from sub-wave-length dimensions in the case of super-resolution imaging techniques to mm if not cm for various light-sheet macroscopes, and no single commercially available test sample covers them all. Advantageously, tissue paper presents structural features spanning several orders of magnitude, making it suitable for many axial sectioning microscopies. **Figs. 3-6** display 3-D images acquired with a large array of techniques.

We first explored the finest structural features with image-scanning microscopy. Also known under its commercial name ‘Airyscan’ microscopy, this technique is a recent variant of confocal laser scanning microscopy (Müller and Enderlein 2010) in which the confocal pinhole is replaced by a honeycomb-type detector array, **Fig. 3***A*. Thus, unlike with the confocal pinhole closed, the detector array collects most of the fluorescence so the signal-to-noise ratio is higher than with conventional confocal detection. Also, the resolution is slightly improved compared with standard confocal microscopy, because the image is reconstructed by reassigning the photons hitting the different detector elements to the ‘right’ image pixel.

On the *z*-stack of images taken on the Airyscan confocal, several tiny features of pyranine-labelled filaments were distinguished, **Fig. 3***B* and **Supplementary Movie S1**. Zooming in on single sections showing these structures in focus revealed protrusions and fibres of sub-wave-length dimensions, with a full width at half maximum (FWHM) of 404 ± 3 nm (*N =* 10, **Fig. 3***C*). Thus, labelled tissue paper contains structural features of dimensions similar to those obtained with fluorescent microspheres (291-320 nm measured FWHM with 93-nm green-fluorescent microspheres, not shown). At the same time, the larger fibres and the surrounding fabric provide valuable context information and permit the assessment of image homogeneity and optical sectioning across the field of view, which in our experiments with the ×63/1.4NA oil objective has a 106-µm image diagonal.

Non-linear microscopy has provided researchers with unique possibilities for biological imaging and photochemistry. It offers attractive advantages, including µm resolution deep in tissue, a largely background-free signal, reduced scattering and better penetration in thick samples, as well as reduced out-of-focus photo damage, which all arise from the square intensity dependence of the excited fluorescence. Non-linear excitation of pyranine (HTPS) fluorescence has been reported at 843 nm (Pastirk, Cruz et al. 2003), whereas ps pulses at 640 nm did not generate appreciable fluorescence in 2P-uncaging experiments (Kiskin, Chillingworth et al. 2002).

We found optimal excitation wavelengths around 820-830 nm and acquired 2P-excited fluorescence images with a ×20/0.8NA air objective on our single-disk ‘virtual light-sheet’ 2PEF spinning-disk microscope, **Fig. 4***A* (Rakotoson, Delhomme et al. 2019). These images reveal a contrasted view of criss-crossing tissue fibres with great detail (400 nm) over a larger FOV than the confocal microscopes (222-µm image diagonal), **Fig. 4***B*. On single optical sections we can distinguish crossing filaments and small features (arrowheads) that are invisible on planes taken only 3 µm apart, illustrating the powerful optical sectioning capacity and versatility of the OA-SIS microscope,

Finally, to illustrate the use of pyranine-labelled tissue paper at larger scales, we used different light-sheet microscopes. The low-fibre density edge of a slanted tissue paper was imaged on a home-built single light sheet SPIM, see **Fig. 5***A*, **Movie S2** and **Fig. S1**. The maximum projection of a *z-*stack of images shows a roughly 1 mm^2^ FOV, **Fig. 5***B*, with an optical sectioning of the order of 25 µm, **Fig. 5***C*. To observe even larger samples areas, we conceived and fabricated on a 3-D printer a small holder to maintain a 2 cm by 2 cm stained tissue paper at a 45° angle with respect to both the illuminating light sheet and detection optical axis, **Fig. 6***A*. We then imaged this large 3-D sample on a light-sheet macroscope at feeble magnification (×1.26/NA0.5). It was difficult to distinguish fine tissue features on single-plane images, **Fig. 6***B*, but the maximum-intensity projection and 3-D rendering clearly reconstructed the slanted 45°-arrangement over a mm height and length scale, **Fig. 6***C*.

Altogether, the images taken on a variety of microscopes show that simple textmarker-labelled tissue paper is commensurable with microscopy techniques spanning the range of hundreds of nm to several mm. Surprisingly small detail can be found with techniques having appropriate sub-wavelength resolution. The relative sparsity of the fibres compared to a fluorescent commercial plastic test slide is advantageous for feature detection while the relative density compared to dispersed-bead samples still allows to assess parameters like the field homogeneity of illumination.

## DISCUSSION

### A simple yet powerful 3-D test sample

This paper proposes a new simple, versatile, easy to use 3-D test sample, well suited for quality control and troubleshooting of fluorescence microscopes. This sample is simply a common tissue paper painted with a “stabilo” pen, in order to stain paper fibres with pyranine dye molecules, a molecule which presents a high photostability and a broad absorption spectrum in blue and UV spectral range. We advocate the routine use of this sample, which is faster, less expensive and complementary to existing calibration procedures.

We show example images for various fluorescence techniques, spanning many orders of magnitude. Our main findings are: (*i*), pyranine, the commonly used fluorophore in such highlighter pens, is very suitable for such calibration purposes, not only in terms of its excitation and emission wavelengths suitable for 488-nm and ‘yellow-green’ detection, but also with respect to its brightness and photostability; (*ii*) at the subwavelength-scale, simple KimWipe^™^ presents near-diffraction limited features, while offering a large, and surprisingly homogeneous and standardised network of fibres from µm up to cm scales; (*iii*) one single swipe results in quite reproducible fluorescent labelling, making the marked tissue paper even useful as a (crude) intensity standard; (*iv*) compatible with a host of different microscopy techniques, the combination of transmission and fluorescence images allows for a direct comparison of microscopes in terms of image contrast and optical sectioning. Our data illustrates the potential of a ‘home-made’, primitive test sample for assessing and comparing day-to-day performance and troubleshooting of high-end microscopes. Its availability, ease of use and virtually zero cost are additional arguments for a routine daily use.

### Images as measurements, measurements from images

Fluorescence measurements gain in quality and reproducibility when a number of parameters are routinely and systematically being monitored,

- the *illumination intensity*, which can vary due to lamp ageing, misalignment, thermal drift, or filter ‘burning’ (i.e., high-intensity damage to their dielectric coatings);
- the illumination and detection *homogeneity*, across the field-of-view;
- the *optical resolution*, which can be degraded due to aberrations, misalignment, a dirty or damaged objective front lens, or even fingerprints on filters;
- the effective axial sectioning obtained in a 3-D sample, which – in addition to the z-resolution also depends on the amount of out-of-focus fluorescence and the density of 3-D labelling, which all impact the contrast in a given focal plane.

Unfortunately, the verification of these parameters is often tedious, labour-intense and typically it requires separate test samples and experiments. Clearly, a simple, inexpensive and multi-scale fluorescent test sample for optical microscopies is needed.

### The quest for an ideal 3-D fluorescent test sample

The combination of specificity, 3-D spatial resolution and the possibility of live-cell time-lapse imaging makes fluorescence microscopy a unique tool for biomedical research (Agard, Hiraoka et al. 1989). Its ability to quantitatively analyse specimens in 3-D allows the inner working of cells, tissues and entire organisms to be probed as never before. Whereas early studies relied on either confocal laser-scanning microscopy (Paddock 1999, Pawley 2006) or wide-field epifluorescence imaging followed by deconvolution (McNally, Karpova et al. 1999, Swedlow 2007), the user nowadays has a bewildering choice among many optical sectioning techniques. This plethora of methods makes it difficult and necessary to evaluate, which of the microscope techniques available in the lab or on an imaging core facility would best respond to a given research question, or if another technique could further improve the 3-D image. David Carters’s chapter 2 in the Paddock book on methods and protocols of confocal microscopy (Paddock 1999) provides a long list of test samples and as many protocols for using such samples for testing instrument performance. A non-exhaustive list includes test for field flatness, axial resolution, chromatic aberrations, z-drive reproducibility, or particle analysis based on identifying objects of different brightness. No less than twelve different samples are listed for such calibration experiments, including large and small beads, silicon chips, slanted mirrors, diatoms, pollen, fluorescent plastic and paper, live cells and plant tissue.

Several flat, 2-D samples are commercially available (like the common fluorescent USAF target or Siemens stars), or they are easily prepared, such as discrete drop-cast beads, thin homogenous dye layers, laser-written fluorescence slides (Corbett, Shaw et all. 2018), aligned macromolecules (Weissman, Klimovsky et al. 2020) or Argolight slides (Royon and Converset 2017). However, the cost for the commonly used beads adds up over the years as they are consumables with a limited shelf life. The Argolight slides on the other hand can be reused, but their initial cost is prohibitively high for many users. In search for a less pricey alternative, Feldhaus and co-workers evaluated in a recent conference poster (Feldhaus, Pan-zera et all. 2019), highlighter pen fluids. They concluded that Zebra 78105 (Mildliner)^™^ and Stabilo Pen 68 neonTM contained spherical sub-resolution fluorescent particles of fairly uniform size that could serve as an alternative to sub-diffraction beads for microscope calibration and performance testing.

Yet, these thin samples are of limited use when it comes to probe 3-D sectioning, along the optical axis. At present, the only de-facto standard for 3-D imaging are the sub-resolution fluorescent microspheres used for PSF measurements, and their embedding in agarose gels can generate fairly flat, yet 3-D samples (Oheim, Salomon et al. 2019). Also, The commercial Argolight calibration slides are designed as calibration standards for troubleshooting, maintenance and alignment. They provide reference fluorescent patterns with stable and precise features (Royon and Converset 2017) but their features in 3-D are small and different lateral scales must be covered by high- and low-magnification variants. On a larger scale, in the range of 1 to 10 micrometers, spiny pollen grains are relatively popular (Potter 1996, Vitha, Bryant et al. 2009, Kirkby, Nadella et al. 2010, Siegel and Brooker 2014, Thériault, Cottet et al. 2014, Kim, Lee et al. 2018, Shin, Kim et al. 2018, Sivaguru, Urban et al. 2018, Rakotoson, Delhomme et al. 2019), and have notably been used for comparison across instruments (Sivaguru, Mander et al. 2012, Sivaguru, Urban et al. 2018). In this context, our textmarker-labeled tissue paper is a cheap, readily available and surprisingly reproducible 3-D multi-scale test sample for day-to-day metrology and troubleshooting.

A different but related problem to day-to-day quality control concerns the choice among different microscope techniques before starting an experiment series, or prior to buying a new instrument. In either case, the question is not to repeatedly monitor a set of parameters over time to ascertain the stability of a measurement, but to identify a “figure-of-merit” and measure it from images of a standardised polyvalent sample taken on several available microscopes. Yet, while the use of images as measurements rather than illustrations is ever more commonplace (Hiemann, Hilger et al. 2006, Waters and Swedlow 2007, Waters 2009, Waters and Wittmann 2014), the road to reproducible fluorescence imaging is less clear, and no consensual metrics have emerged, despite several efforts of standardisation (Model and Burkhardt 2001, Brown, Reilly et al. 2015, Deagle, Wee et al. 2017, Alexia, Schleicher et al. 2019). Several studies have pointed out the need for validation and reproducibility in the acquisition (Murray, Appleton et al. 2007) and image treatment of 3-D data sets (Dieterlen, Xu et al. 2002). With our pyranine-labelled tissue sample, we provide an effective solution for these applications, at minimal cost.

Without wanting to replace existing and proven tools and procedures, our test sample offers a complementary, quick and straightforward possibility to assess instrument performance and monitor several important image features from a single image or image stack. We believe it to be an ideal sample for prototyping, routine quality assessment and rapid troubleshooting during an ongoing experiment.

### Pyranine as a fluorescence standard

Pyranine is an inexpensive, hydrophilic, pH-sensitive fluorescent dye from the group of aryl-sulfonates. The molecule has applications as a colouring agent, biological stain, optical detection reagent, and as a pH indicator with a pKa of ∼7.3 in aqueous buffers, and it has been used as a stromal and cameral pH sensor *in vivo* (Thomas, Brimijoin et al. 1990). It is impermeable to biological membranes and non-toxic under all realistic concentrations, making it a suitable tracer and parent fluorophore for various derivatives in live-cell applications (see, e.g. (Bort, Gallavardin et al. 2013, Legenzov, Dirda et al. 2015)), including mixed 1P-imaging and 2P-uncaging experiments (Kiskin, Chillingworth et al. 2002). Pyranine also acts as a fluorescent chemosensor (in the green emission band) for copper, even in a competitive environment, with a 2:1 stoichiometry (Saha, Sengupta et al. 2014). Our measured excitation and emission spectra (Fig.2, Fig.S2) clearly suggest that the fluorophore used in yellow highlighter pen is pyranine. Our identification of pyranine is corroborated by several online resources^1^.

## CONCLUSION

We here extended the work of Feldhaus and co-workers (2019) to a volumetric, 3-D sample for optical sectioning microscopies. There is some truth to the old saying “simple yet effective”, at least when it comes to highlighter pen-labelled tissue paper as a home-made 3-D test sample. With spatial features covering several orders of magnitude, a random but fairly reproducible

fibre density and fluorescence intensity and the ‘biology-like’ aspect are clear pluses. The simple preparation in a minute and virtually zero cost make it even more attractive. Many more colour variants in addition to yellow are available and many different fabrics can be used, making our procedure a versatile approach for standardisation and quality control in quantitative fluorescence microscopy.

### BOX1.

*3-D deconvolution microscopy* or ‘exhaustive photon reassignment’ is a computational de-blurring method used to reduce the unwanted out-of-focus fluorescence in 3-D microscope images. It is compatible with (but not restricted to) epifuorescence excitation, and – based on the precise 3-D optical response of the microscope optics - it puts photons back into the plane where the originated from. The 3-D image is considered as a convolution of the true fluorophore distribution with the microscope optics’ point-spread function (PSF). With knowledge of the PSF, the true fluorophore distribution can be back-calculated from a 3-D image stack by numerical de-convolution. However, the technique is inherently noise-sensitive and thus requires a good signal-to-noise ratio. Also, it is only as good as the knowledge of the PSF, which can vary across the sample volume. In fact deconvolution artefacts are so notorious that it has become good practice to use sub-diffraction fluorescent microspheres embedded in an agarose gel as a 3-D test sample and to convince oneself of the plausibility of the reconstructed image prior to studying 3-D biological samples of unknown fluorophore distributions.

This work contains **Supplementary material**, online.

## ACKNOWLEDGEMENTS

Spectral and Airyscan confocal micrographs were taken on microscopes of the imaging core facility (CNRS UMS 2009, INSERM US 36, BioMedTech Facilities). Marcoscopic light-sheet images were acquired at UtechS Photonic BioImaging (Imagopole), C2RT, Institut Pasteur, supported by the French National Research Agency (France BioImaging; ANR-10–INSB–04; Investments for the Future). AS acknowledges support from the French Ministry of Foreign Affairs and the French Embassy in Tel Aviv (Chateaubriand fellowship). KS is the laureate of a joint CNRS-U Toronto PhD fellowship. This study was financed by the CNRS, the University of Paris, the Israeli Science Foundation (ISF, grant 1231/19), U Toronto and the European Union (H2020 Eureka! Eurostars, ‘NanoScale’, E!12848, https://nanoscale.sppin.fr, to MO & S).

The authors are grateful for mobility support from a Franco-Israeli CNRS-LIA, ‘ImagiNano’, and the FranceBioImaging large-scale national infrastructure initiative (FBI, ANR-10-INSB-04, Investments for the future). These funding agencies had no role in study design, data collection and analysis, decision to publish, or preparation of the manuscript.

## Footnotes

1 https://www.compoundchem.com/2015/01/22/highlighters/, https://en.wikipedia.org/wiki/Highlighter; https://en.wikipedia.org/wiki/Pyra-nine; https://www.chemeurope.com/en/infographics/177/the-chemistry-of-highlighter-colours.html;

## REFERENCES

Agard, D. A., Y. Hiraoka, P. Shaw and J. W. Sedat (1989). “Fluorescence microscopy in three dimensions.” Methods in cell biology 30: 353–377.

Alexia, F., K. D. Schleicher, E. Nikolaus, H. Wolf and B. Oliver (2019). “Using the NoiSee workflow to measure signal-to-noise ratios of confocal microscopes.” Scientific Reports (Nature Publisher Group) 9(1).

Barentine, A. E., L. K. Schroeder, M. Graff, D. Baddeley and J. Bewersdorf (2018). “Simultaneously measuring image features and resolution in live-cell STED images.” Biophysical journal 115(6): 951–956.

Bort, G., T. Gallavardin, D. Ogden and P. I. Dalko (2013). “From one photon to two photon probes:”caged” compounds, actuators, and photoswitches.” Angewandte Chemie International Edition 52(17): 4526–4537.

Brown, C. M., A. Reilly and R. W. Cole (2015). “A quantitative measure of field illumination.” Journal of biomolecular techniques: JBT 26(2): 37.

Corbett, A. D., M. Shaw, A. Yacoot, A. Jefferson, et al. (2018) Microscope calibration using laser written fluorescence. Optics Express 26(17): 21887–900

De Borba, E., C. Amaral, M. Politi, R. Villalobos and M. Baptista (2000). “Photophysical and photochemical properties of pyranine/methyl viologen complexes in solution and in supramolecular aggregates: A switchable complex.” Langmuir 16(14): 5900–5907.

Deagle, R. C., T.-L. E. Wee and C. M. Brown (2017). “Reproducibility in light microscopy: Maintenance, standards and SOPs.” The international journal of biochemistry & cell biology 89: 120–124.

Denk, W., J. H. Strickler and W. W. Webb (1990). “Two-photon laser scanning fluorescence microscopy.” Science 248(4951): 73–76.

Diaspro, A. (2002). Confocal and two-photon microscopy: foundations, applications, and advances, Wiley-Liss New York:.

Dieterlen, A., C. Xu, M.-P. Gramain, O. Haeberlé, B. Colicchio, C. Cudel, S. Jacquey, E. Ginglinger, G. Jung and É. Jeandidier (2002). “Validation of image processing tools for 3-D fluorescence microscopy.” Comptes rendus biologies 325(4): 327–334.

Dodt, H.-U., U. Leischner, A. Schierloh, N. Jährling, C. P. Mauch, K. Deininger, J. M. Deussing, M. Eder, W. Zieglgänsberger and K. Becker (2007). “Ultramicroscopy: three-dimensional visualization of neuronal networks in the whole mouse brain.” Nature methods 4(4): 331–336.

Feldhaus, C., A. Panzera, R. Palmisano. (2019) “Evaluation of highlighter pens as cheap and cheerful samples for microscope calibration and performance testing. Conference presentation/poster. Focus on microscopy (FOM), 2019, London. See https://webdav.tuebingen.mpg.de/LM/FOM2019/ for more infos.

Hiemann, R., N. Hilger, U. Sack and M. Weigert (2006). “Objective quality evaluation of fluorescence images to optimize automatic image acquisition.” Cytometry Part A: The Journal of the International Society for Analytical Cytology 69(3): 182–184.

Hillman, E. M., V. Voleti, W. Li and H. Yu (2019). “Light-sheet microscopy in neuroscience.” Annual review of neuroscience 42: 295–313.

Huber, D., M. Keller and D. Robert (2001). “3D light scanning macrography.” Journal of microscopy 203(2): 208–213.

Huisken, J., J. Swoger, F. Del Bene, J. Wittbrodt and E. H. Stelzer (2004). “Optical sectioning deep inside live embryos by selective plane illumination microscopy.” Science 305(5686): 1007–1009.

Keller, P. J. and H.-U. Dodt (2012). “Light sheet microscopy of living or cleared specimens.” Current opinion in neurobiology 22(1): 138–143.

Kim, G., S. Lee, S. Shin and Y. Park (2018). “Three-dimensional label-free imaging and analysis of Pinus pollen grains using optical diffraction tomography.” Scientific reports 8(1): 1–8.

Kirkby, P. A., K. N. S. Nadella and R. A. Silver (2010). “A compact acousto-optic lens for 2D and 3D femtosecond based 2-photon microscopy.” Optics express 18(13): 13720–13744.

Kiskin, N. I., R. Chillingworth, J. A. McCray, D. Piston and D. Ogden (2002). “The efficiency of two-photon photolysis of a” caged” fluorophore, o-1-(2-nitrophenyl) ethylpyranine, in relation to photodamage of synaptic terminals.” European Biophysics Journal 30(8): 588–604.

Legenzov, E. A., N. D. Dirda, B. M. Hagen and J. P. Kao (2015). “Synthesis and characterization of 8-O-carboxymethylpyranine (CM-pyranine) as a bright, violet-emitting, fluid-phase fluorescent marker in cell biology.” PloS one 10(7): e0133518.

Masters, B. R. (2020). Richard Zsigmondy and Henry Siedentopf’s Ultramicroscope. Superresolution Optical Microscopy, Springer: 165–172.

McNally, J. G., T. Karpova, J. Cooper and J. A. Conchello (1999). “Three-dimensional imaging by deconvolution microscopy.” Methods 19(3): 373–385.

Model, M. A. and J. K. Burkhardt (2001). “A standard for calibration and shading correction of a fluorescence microscope.” Cytometry: The Journal of the International Society for Analytical Cytology 44(4): 309–316.

Müller, C. B. and J. Enderlein (2010). “Image scanning microscopy.” Physical review letters 104(19): 198101.

Murray, J. M., P. L. Appleton, J. R. Swedlow and J. C. Waters (2007). “Evaluating performance in three dimensional fluorescence microscopy.” Journal of microscopy 228(3): 390–405.

Nadrigny, F., I. Rivals, P. G. Hirrlinger, A. Koulakoff, L. Personnaz, M. Vernet, M. Allioux, M. Chaumeil, N. Ropert and C. Giaume (2006). “Detecting fluorescent protein expression and co-localisation on single secretory vesicles with linear spectral unmixing.” European Biophysics Journal 35(6): 533–547.

Neher, E. (1995). “The use of fura-2 for estimating Ca buffers and Ca fluxes.” Neuropharmacology 34(11): 1423–1442.

Oheim, M., A. Salomon, A. Weissman, M. Brunstein and U. Becherer (2019). “Calibrating evanescent-wave penetration depths for biological TIRF microscopy.” Biophysical Journal 117(5): 795–809.

Paddock, S. W. (1999). Confocal microscopy: methods and protocols, Springer Science & Business Media.

Pastirk, I., J. M. D. Cruz, K. A. Walowicz, V. V. Lozovoy and M. Dantus (2003). “Selective two-photon microscopy with shaped femtosecond pulses.” Optics express 11(14): 1695–1701.

Pawley, J. (2006). Handbook of biological confocal microscopy, Springer Science & Business Media.

Potter, S. M. (1996). “Vital imaging: two photons are better than one.” Current Biology 6(12): 1595–1598.

Rakotoson, I., B. Delhomme, P. Djian, A. Deeg, M. Brunstein, C. Seebacher, R. Uhl, C. Ricard and M. Oheim (2019). “Fast 3-D imaging of brain organoids with a new single-objective planar-illumination two-photon microscope.” Frontiers in neuroanatomy 13: 77.

Resch-Genger, U., K. Hoffmann, W. Nietfeld, A. Engel, J. a. Neukammer, R. Nitschke, B. Ebert and R. Macdonald (2005). “How to improve quality assurance in fluorometry: fluorescence-inherent sources of error and suited fluorescence standards.” Journal of Fluorescence 15(3): 337–362.

Royon, A. and N. Converset (2017). “Quality Control of Fluorescence Imaging Systems: A new tool for performance assessment and monitoring.” Optik & Photonik 12(2): 22–25.

Saha, T., A. Sengupta, P. Hazra and P. Talukdar (2014). “In vitro sensing of Cu+ through a green fluorescence rise of pyranine.” Photochemical & Photobiological Sciences 13(10): 1427–1433.

Schindelin, J., I. Arganda-Carreras, E. Frise, V. Kaynig, M. Longair, T. Pietzsch, S. Preibisch, C. Rueden, S. Saalfeld and B. Schmid (2012). “Fiji: an open-source platform for biological-image analysis.” Nature methods 9(7): 676–682.

Shin, Y., D. Kim and H. S. Kwon (2018). “Oblique scanning 2 photon light sheet fluorescence microscopy for rapid volumetric imaging.” Journal of biophotonics 11(5): e201700270.

Siedentopf, H. and R. Zsigmondy (1902). “Uber sichtbarmachung und größenbestimmung ultramikoskopischer teilchen, mit besonderer anwendung auf goldrubingläser.” Annalen der Physik 315(1): 1–39.

Siegel, N. and G. Brooker (2014). “Improved axial resolution of FINCH fluorescence microscopy when combined with spinning disk confocal microscopy.” Optics express 22(19): 22298–22307.

Sivaguru, M., L. Mander, G. Fried and S. W. Punyasena (2012). “Capturing the surface texture and shape of pollen: a comparison of microscopy techniques.” PloS one 7(6): e39129.

Sivaguru, M., M. A. Urban, G. Fried, C. J. Wesseln, L. Mander and S. W. Punyasena (2018). “Comparative performance of airyscan and structured illumination superresolution microscopy in the study of the surface texture and 3D shape of pollen.” Microscopy research and technique 81(2): 101–114.

Swedlow, J.R. (2007). “Quantitative fluorescence microscopy and image deconvolution.” Methods in cell biology 81: 447–465.

Taniguchi, M. and J. S. Lindsey (2018). “Database of absorption and fluorescence spectra of> 300 common compounds for use in photochem CAD.” Photochemistry and photobiology 94(2): 290–327.

Thériault, G., M. Cottet, A. Castonguay, N. McCarthy and Y. De Koninck (2014). “Extended two-photon microscopy in live samples with Bessel beams: steadier focus, faster volume scans, and simpler stereoscopic imaging.” Frontiers in cellular neuroscience 8: 139.

Thomas, J. V., M. R. Brimijoin, T. R. Neault and R. F. Brubaker (1990). “The fluorescent indicator pyranine is suitable for measuring stromal and cameral pH in vivo.” Experimental eye research 50(3): 241–249.

van’t Hoff, M., M. Reuter, D. T. Dryden and M. Oheim (2009). “Screening by imaging: scaling up single-DNA-molecule analysis with a novel parabolic VA-TIRF reflector and noise-reduction techniques.” Physical Chemistry Chemical Physics 11(35): 7713–7720.

Vitha, S., V. M. Bryant, A. Zwa and A. Holzenburg (2009). “3D confocal imaging of pollen.” Microscopy and Microanalysis 15(S2): 622–623.

Voie, A. H., D. Burns and F. Spelman (1993). “Orthogonal plane fluorescence optical sectioning: Three dimensional imaging of macroscopic biological specimens.” J. Microsc. 170(3): 229–236.

Wan, Y., K. McDole and P. J. Keller (2019). “Light-sheet microscopy and its potential for understanding developmental processes.” Annual review of cell and developmental biology 35: 655–681.

Waters, J. C. (2009). Accuracy and precision in quantitative fluorescence microscopy, The Rockefeller University Press.

Waters, J. C. and J. R. Swedlow (2007). “Interpreting fluorescence microscopy images and measurements.” Evaluating Techniques in Biochemical Research. D. Zuk, editor. Cell Press, Cambridge, MA: 37–42.

Waters, J. C. and T. Wittmann (2014). Concepts in quantitative fluorescence microscopy. Methods in cell biology, Elsevier. 123: 1–18.

Weissman, A., H. Klimovsky, D. Harel, R. Ron, M. Oheim and A. Salomon (2020). “Fabrication of dipole-aligned thin films of Porphyrin J-aggregates over large surfaces.” Langmuir 36(4): 844–851.

Zhang, B., J. Zerubia and J.-C. Olivo-Marin (2007). “Gaussian approximations of fluorescence microscope pointspread function models.” Applied optics 46(10): 1819–1829.

