## SUPPLEMENTARY MATERIAL for "A simple, inexpensive and multi-scale 3-D fluorescent test sample for optical sectioning microscopies"

*Short title*: 3-D fluorescent test sample

Ilya Olevsko,[1,*] Kaitlin Szederkenyi,[2,3,*] Jennifer Corridon,[4,5] Aaron Au,[3] Brigitte Delhomme,[2] Thierry Bastien, [4,6] Julien Fernandes,[7] Christopher Yip,[3] Martin Oheim,[2,4,5] and Adi Salomon,[1,2,a*,] 🖂

[1] Institute of Nanotechnology and Advanced Materials (BINA), Department of Chemistry, Bar-Ilan University, Ramat-Gan, 52900 Israel;
[2] Université de Paris, CNRS, SPPIN - Saints-Pères Paris Institute for the Neurosciences, F-75006 Paris, France;
[3] University of Toronto, Donnelly Centre for Cellular & Biomolecular Research, 160 College St, Toronto, ON M5S 3E1, Canada;
[4] Université de Paris, CNRS UMS 2009, INSERM US 36, BioMedTech Facilities, Campus Saint Germain, F-76006 Paris, France;
[5] SCM – *Service Commun de Microscopie*, Université de Paris, Campus Saint Germain, F-76006 Paris, France;
[6]*Plateforme de Prototypage*, Université de Paris, Campus Saint Germain, F-76006 Paris, France;
[7]Institut Pasteur, UTechS Photonic BioImaging C2RT, F-75015 Paris.

[#]Co-first authors

This supplementary material contains

- A list of abbreviations p. 2
- Table S1 p. 3
- Supplementary Figures S1 – S2 p. 4
- Supporting Movies S1 – S2 p. 6

#### List of used abbreviations

2P - two-photon (excitation fluorescence)

| | | |
|---|---|---|
| 3-D | - | three-dimensional |
| AU | - | arbitrary units |
| BU | - | bead units |
| sCMOS | - | scientific complementary metal-oxyde semiconductor |
| CNRS | - | Centre National de la Recherche Scientifique |
| CWL | - | center wavelength |
| CLSM | - | confocal laser scanning microscopy |
| EGFP | - | enhanced green-fluorescent protein |
| EMCCD | - | electron-multiplying charge-coupled device |
| EYPF | - | enhanced yellow-fluorescent protein |
| FITC | - | fluorescein isothiocyanate a |
| FOV | - | field of view |
| FWHM | - | full width at half maximum |
| HPTS | - | 8-hydroxypyrene-1,3,6-trisulfonic acid trisodium salt |
| LP | - | long pass (filter) |
| LSM | - | laser-scanning microscope |
| LUT | - | look-up table |
| NA | - | numerical aperture |
| OASIS | - | On-Axis 2-Photon Light-Sheet Imaging System |
| PALM | - | photo-activation localisation microscopy |
| PSF | - | point-spread function |
| SAF | - | supercritical angle fluorescence |
| SD | - | standard deviation |
| SPIM | - | selective-plane illumination microscopy |
| STORM | - | stochastic optical reconstruction microscopy |
| TIRF | - | total internal reflection fluorescence |
| TIRTC | - | tetramethylrhodamine isothiocyanate |
| USAF | - | United States Air Force |
| UV | - | ultraviolet |

### Supplementary Table

**Table S1**

*Properties of the used filter sets*

| | ex. | dic | em | *Ref., comments* |
|---|---|---|---|---|
| Epifluorescence | | | | |
| Fura-2 HC | 387/11 BrightLine | 409 BS | 510/84 BrightLine | AHF F76-520 |
| TRITC HC | 543/22 BrightLine | 562 BS | 593/40 BrightLine | AHF F36-503 |
| Confocal LSM710 | | | | |
| 488-nm $Ar^+$ line | - | 488 HFT | 510-700 | META detector |
| Airyscan LSM880 | | | | |
| 488-nm $Ar^+$ line | - | 488 MFT | BP495-550+LP580 | dual band emission |
| OASIS 2P-sppinging disk | | | | |
| 820 nm Ti:Sapph* | - | 715SP + 705SP + 672SP + 565SP | ET700SP-2P + 400-565 | only green channel used |
| Home-built SPIM (light sheet) | | | | |
| 488 nm DPPS | - | 500LP | 553/44BP | 90° geometry |
| Ultramicroscope II (light sheet) | | | | |
| White laser | 470/40X | - | 525/50M | 90° geometry |

* *two-photon excitation*; BP – band-pass; ET- enhanced transmission; HC – hard coated; HFT – main dichroic ("*Hauptfarbteiler*" in German); MFT – multi-line dichroic.

### Supplementary Figures

*Home-built SPPIM*

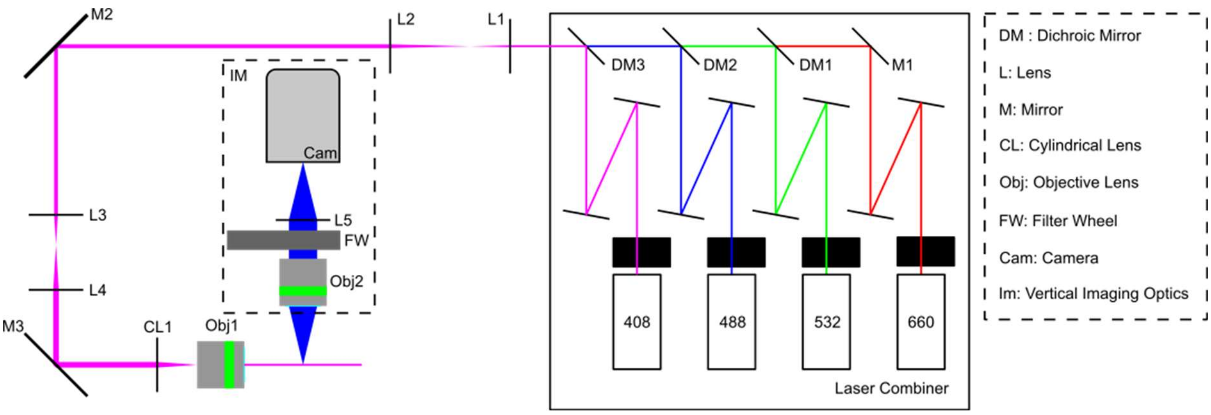


**Figure S1.** *Simplified optical layout of the home-built SPIM.* Schematic optical layout of our modular light-sheet microscope built from optical bench components. Four lasers (408 nm, Coherent Cube, 50 mW; 488 nm, Coherent Sapphire, 20 mW; 532 nm, Laser Quantum GEM, 400 mW; 660 nm, Laser Quantum IGNIS, 400 mW) were combined, the beam expanded 6.66-fold with a first ×2 ($f_1$ = 30 mm and $f_2$ = 60 mm achromatic (AC) doublets) and second ×3.33 telescope (AC $f_3$ = 30 mm and $f_2$ = 100 mm) and focused to a light sheet by a plano-convex cylindrical lens ($f_{CL1}$ = 40 mm, Thorlabs LJ1402L1-A). Both the illuminating and imaging objectives were Olympus SLMPlan ×20/0.35NA, the tube lens a Thorlabs AC $f_5$ = 150 mm. The camera was an Andor NEO 5.5. sCMOS.

*Identification of the fluorophore as pyranine*

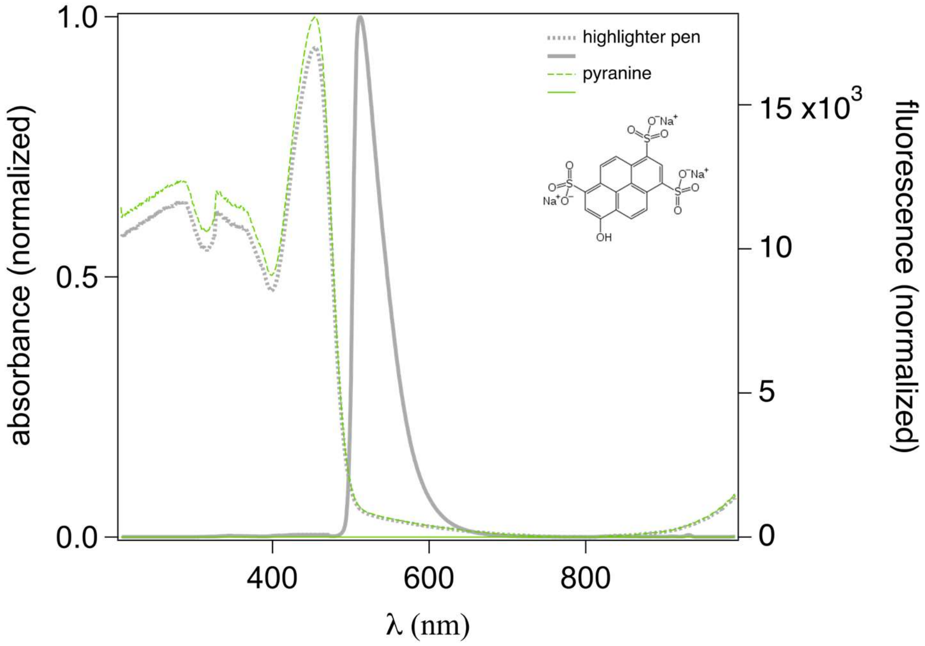


**Figure S3**. *Comparison of measured highlighter pen dye (grey) and pyranine spectra (green).* Pyranine, 8-Hydroxypyrene-1,3,6-trisulfonic acid, HPTS ($C_{16}H_7Na_3O_{10}S_3$, also known as Solvent Green 7; sulfonated hydroxy pyrene trisodium salt, CAS 6358-69-6, see *inset* for its chemical structure) is a yellow-green crystalline powder soluble in water and in various alcohols. We dissolved pyranine to ~10 µM in a 20% EtOH/water mixture and measured the absorption from a thin liquid film (broken line). To this end, 20 µl of solution were sandwiched between two microscope coverslips. Fluorescence (through line) was measured from the same sample upon 450-nm excitation through a 500LP emission filter and 490LP dichroic mirror. The graph also reproduces, for a direct comparison, the data from fig.2, which was measured from a smear of highlighter on a clean microscope slide. Fluorescence filter sets were identical for both dyes. The tiny differences between the absorption spectra can be attributed to solvent effects and different pH. Our identification of pyranine is corroborated by Feldhaus et al. (2019), among others.

### Supplementary Movies

**<u>Movie S1</u>**

*3-D rendered z-scan with the Airyscan confocal microscope*

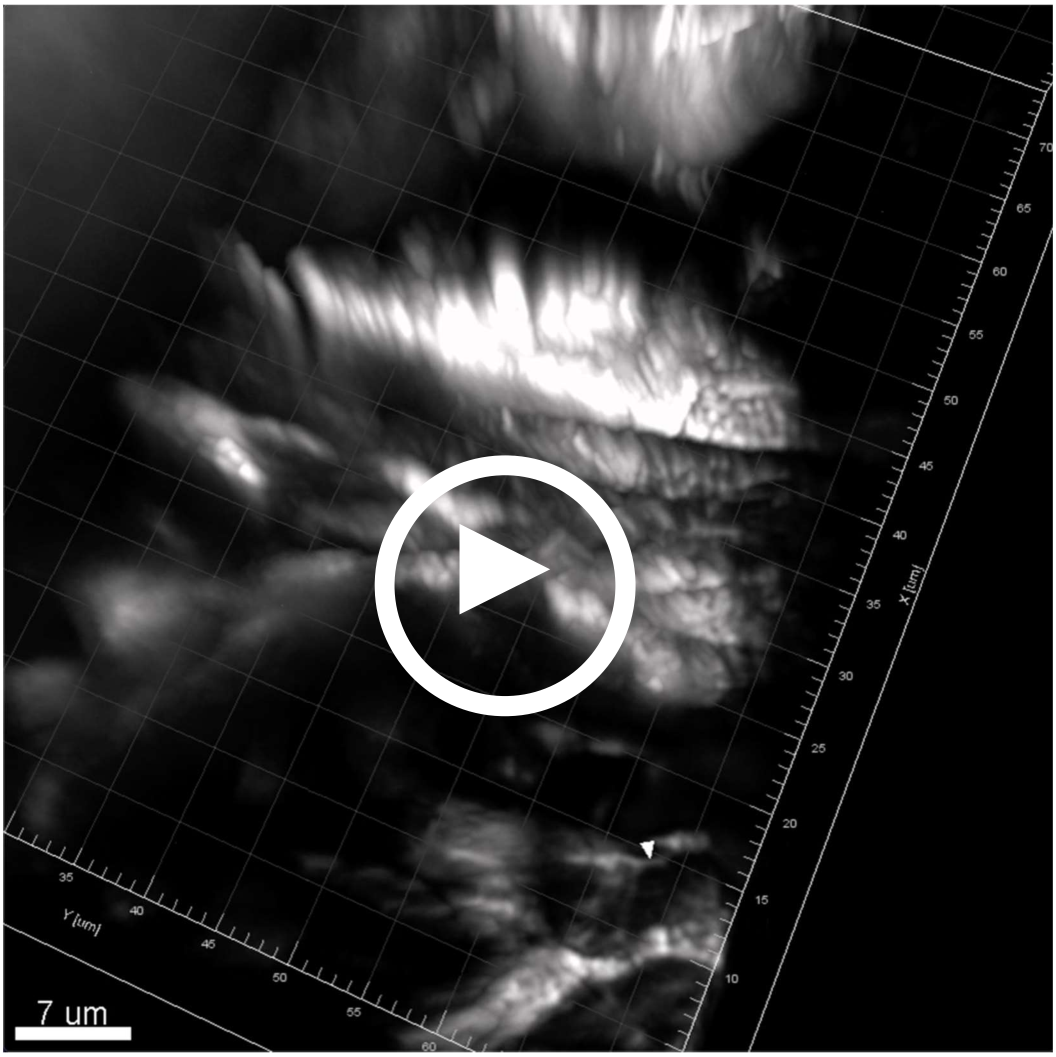


**Movie S1.** *Axial scan with the Airyscan confocal microscope.* 3-D Reconstruction of a *z*-stack of fluorescence images of a KimWipe® stripe, labelled with a yellow Stabilo® highlighter pen. 107 consecutive optical slices were acquired on a Zeiss LSM 880 Fast Airyscan confocal microscope (Plan Apochromat ×63/NA 1.4oil). We used an axial step size of 187 nm during acquisitions, the *z*-stack reconstructs a total of almost 20 µm (19.98 µm). The arrow head points at a structure having a measured FWHM of 383 nm. Pixel size was 43 nm in the sample plane, image size 73.95 × 73.95 µm² - Speed of animation of the 3D reconstruction: 24 fps. See MRT journal website for link to the video file.

**Movie S2**

*Axial scan with the home-built light-sheet microscope*

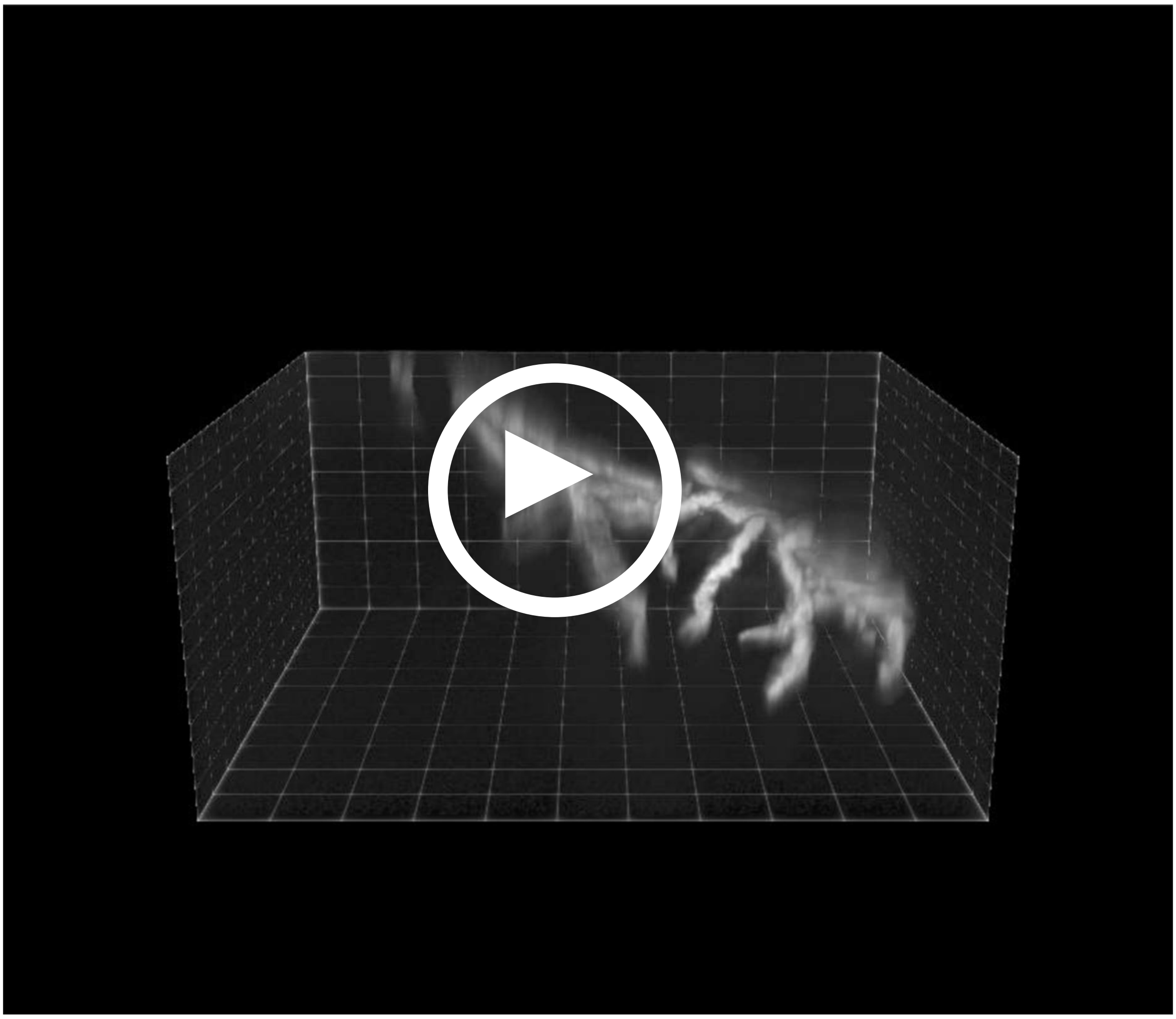

**Movie S2.** *Axial scan with the home-built light-sheet microscope. z*-stack from the fibrous border of a KimWipe® stripe, labelled with a yellow highlighter pen. 470 slices were taken on a custom light sheet microscope. Objectives were both SLMPlan 20x/0.35. We used a step size of 1 µm during acquisitions, the *z*-stack reconstructs a total of 470 µm. The effective pixel size was 39 nm in the sample plane, image size 998.4 µm x 842.4 µm – Speed of animation of the 3D reconstruction: 14 fps. See MRT journal website for link.
